# Regulation of proteostasis by sleep through autophagy in *Drosophila* models of Alzheimer’s Disease

**DOI:** 10.1101/2024.07.01.601554

**Authors:** Natalie Ortiz-Vega, Amanda G. Lobato, Tijana Canic, Yi Zhu, Stanislav Lazopulo, Sheyum Syed, R. Grace Zhai

## Abstract

Sleep and circadian rhythm dysfunctions are common clinical features of Alzheimer’s Disease (AD). Increasing evidence suggests that in addition to being a symptom, sleep disturbances can also drive the progression of neurodegeneration. Protein aggregation is a pathological hallmark of AD, however the molecular pathways behind how sleep affects protein homeostasis remain elusive. Here we demonstrate that sleep modulation influences proteostasis and the progression of neurodegeneration in *Drosophila* models of Tauopathy. We show that sleep deprivation enhanced Tau aggregational toxicity resulting in exacerbated synaptic degeneration. In contrast, sleep induction using gaboxadol led to reduced hyperphosphorylated Tau accumulation in neurons as a result of modulated autophagic flux and enhanced clearance of ubiquitinated Tau, suggesting altered protein processing and clearance that resulted in improved synaptic integrity and function. These findings highlight the complex relationship between sleep and autophagy, in regulating protein homeostasis, and the neuroprotective potential of sleep-enhancing therapeutics to slow the progression or delay the onset of neurodegeneration.

## Introduction

Alzheimer’s disease (AD) is the most common neurodegenerative disorder and the leading cause of dementia [1]. Affected individuals display a variety of progressive and disabling neurological symptoms such as cognitive decline, motor dysfunction, and psychiatric symptoms [2]. The main pathological phenotype of Alzheimer’s disease is shrinkage of the hippocampus and cortex, and the accumulation of plaques formed by extracellular insoluble aggregates of amyloid-beta fragments (Aβ), and neurofibrillary tangles (NFTs) formed by intracellular accumulation of hyperphosphorylated Tau (pTau) [3, 4]. Tau proteins are microtubule binding proteins that promote stabilization and assembly of the microtubules. Under pathological conditions, Tau is hyperphosphorylated, detached from microtubules, and form neurotoxic oligomers that eventually aggregate into paired helical fragments (PHF) and eventually form NFTs [5]. In AD brains, Tau is three to four-fold more hyperphosphorylated than normal adults, causing disruption of its normal function and facilitating its polymerization into paired helical fragments (PHF) and forming NFTs [6].

Sleep disturbances are observed in approximately 45% of patients and may be one of the first manifestations of the disease [2, 7, 8]. These sleep disorders include insomnia, excessive daytime sleepiness (EDS), and fragmented sleep [2]. Specifically, polysomnography studies show reduced total sleep time and sleep efficiency, increased sleep-onset latency, and wake time after sleep onset, reduced deep sleep an REM sleep amounts [2, 9]. This is particularly important because increasing evidence suggests a bidirectional relationship between sleep and neurodegenerative diseases, where on one side, neurodegeneration can lead to circadian dysregulation and sleep disorders [10, 11]; While on the other hand, sleep disruption can promote neurodegeneration through induced neuroinflammation, and aberrant protein homeostasis [12]. Meta-analysis results suggest that individuals with disturbed sleep have a 1.68 times higher risk to develop cognitive impairment and/or AD [13]. Moreover, sleep deprivation is shown to affect the glymphatic clearance of proteins in the brain and increases the accumulation of Aβ in the hippocampal region, and the accumulation of hyperphosphorylated Tau leading to inhibition of neurogenesis and cognitive dysfunction [13–16].

Recent studies suggest that sleep regulates autophagy in a daily sleep/wake cycle, and sustained changes in autophagosome level affects sleep amount [17]. Although sleep disorders have been linked to multiple neurodegenerative diseases, few studies have fully investigated how sleep affects AD progression. Previous studies have indicated that sleep abnormalities exacerbate neurodegeneration and that sleep therapies may slow down disease progression. For example, in a transgenic mouse model of Aβ deposition, it has been observed that sleep deprivation accelerates plaque accumulation, whereas promoting sleep with an orexin antagonist significantly inhibits plaque formation [18]. However, the molecular mechanisms of how these interventions lead to pathophysiological changes in AD remain unclear.

Sleep abnormalities are highly present in AD patients prior to clinical onset of the disease [19], therefore, uncovering the sleep-neurodegeneration relationship longitudinally and prior to disease manifestation is required. *Drosophila* is a powerful genetic model system to study AD and sleep [20] Several models have been characterized to model given its complex etiology [21, 22]. In this report, we took advantage of well characterized *Drosophila* models of Tauopathy to address the molecular mechanisms of how sleep abnormalities affect proteostasis and Tau-induced neurodegeneration *in vivo*.

## Results

### Establish a sleep modulation paradigm to assess altered progression of neurodegeneration

Similar to humans, *Drosophila* follows a 12-hour day/12-hour night rhythm [23]. Sleep in flies is measured as 5 minutes or more of inactivity [24, 25]. To study *Drosophila* locomotor activity and sleep we used a *Drosophila* Activity Monitor (TriKinetics, Waltham, MA) using a locally built detection system that includes and infra-red (IR) beam to record locomotor activity of solitary flies placed in glass tubes **(Figure S1A-B)**, as commonly used in measuring fly sleep [26]. Using the activity monitoring system, we first determined the sleep patterns of Tauopathy models. *Drosophila* models for tauopathy, established by expressing either wild type (Tau^WT^) or mutant (Tau^R406W^) human Tau protein in neurons, recapitulate adult-onset, progressive neurodegeneration with altered lifespan and accumulation of pTau [27]. We expressed either *UAS-CD8-GFP*, *UAS-Tau^WT^* or *UAS-Tau^R406W^* pan-neuronally using *elav-GAL4* driver or in the photoreceptors using *GMR-GAL4* driver as previously reported [27, 28], and monitored *Drosophila* activity behavior for 7 days, which included 2 days of acclimation and 4 days of sleep monitoring (**Figure 1A**).

**Figure 1:**
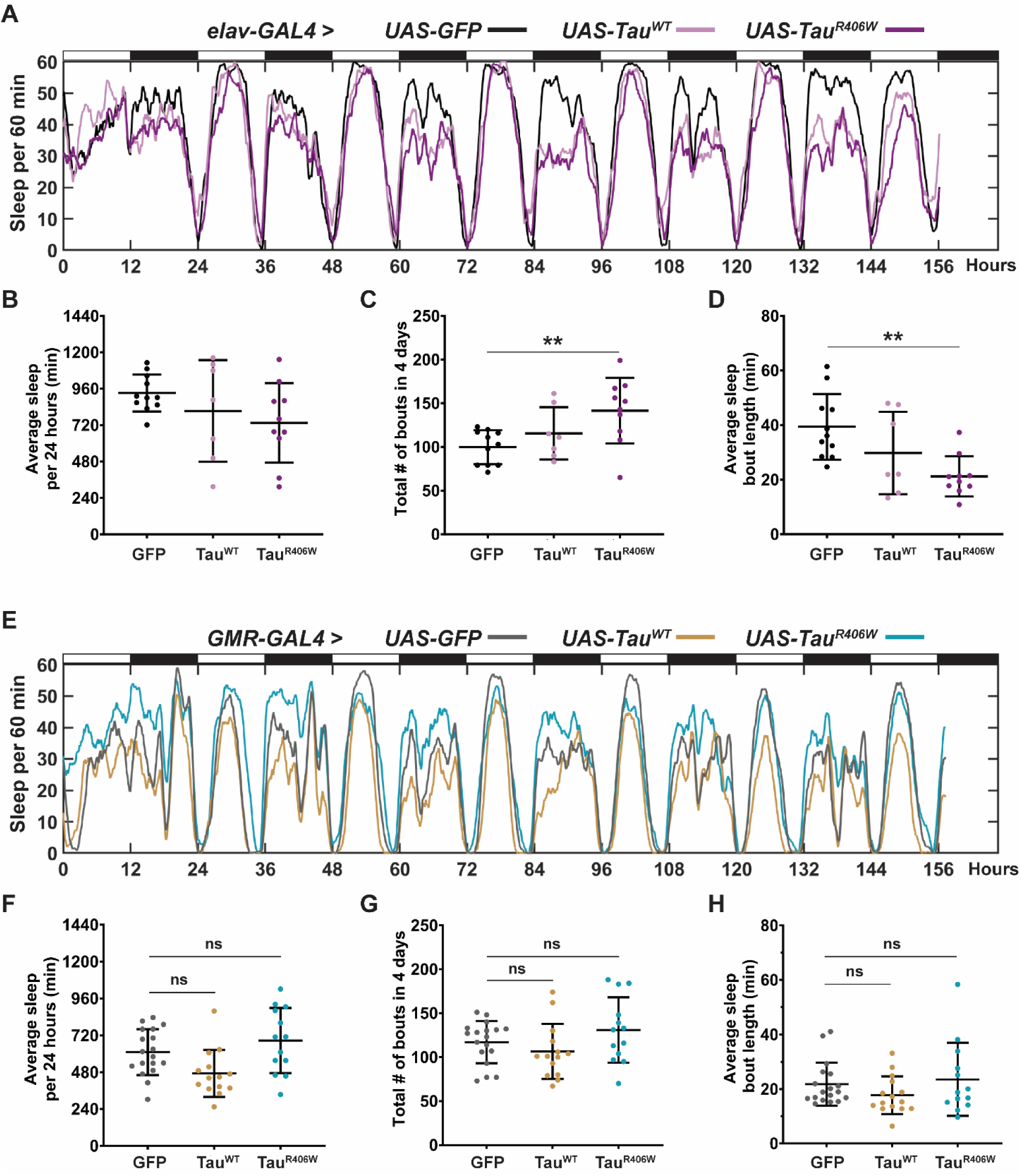
Photoreceptor expression of hTau causes minimum effects on sleep. **(A)** Sleep profiles of 2 DAE flies expressing either *UAS-CD8-GFP* (black), *UAS-hTau^WT^* (light pink) or *UAS-hTau^R406W^* (magenta^)^ pan-neuronally using *elav-GAL4* driver. Flies were allowed to acclimate from 0-48 hours and sleep was measured from 48-144 hours. **(B)** Quantification of average sleep per 24 hours measured in minutes. **(C)** Quantification of total number of bouts in 4 days (40-144 hours). **(D)** Quantification of average sleep bout length in minutes. **(E)** Sleep profiles of 2 DAE flies expressing either *UAS-CD8-GFP* (gray), *UAS-hTau^WT^* (gold) or *UAS-hTau^R406W^* (turquoise) in the photoreceptors using *GMR-GAL4* driver. **(F)** Quantification of average sleep per 24 hours measured in minutes. **(G)** Quantification of total number of bouts in 4 days. **(H)** Quantification of average sleep bout length in minutes. Data as mean ± SD. n = 7-22, One-way ANOVA, *\*\*p<0.01*.

Pan-neuronal expression of Tau^WT^ or Tau^R406W^ with *elav-GAL4* caused significant sleep behavior changes, specifically in reduced average sleep time per 24 hours of 117 min less in Tau^WT^, and 197 min less in Tau^R406W^ (**Figure 1B**); increased number of sleep bouts (**Figure 1C**), and decreased length of each sleep bout **(Figure 1D)**. Combined, these results show increased sleep fragmentation and decreased total sleep length, suggesting that Tau^WT^ and Tau^R406W^ expression in all neurons causes sleep disturbances, consistent with previous reports that have observed sleep disturbances caused by pan-neuronal expression of Aβ or Tau [29–31]. Such significant change in sleep behavior makes it challenging to dissect the direct effects of sleep modulation on neurodegeneration. We therefore sought to identify a more suitable tauopathy model by expressing Tau in a subset of neurons with minimum impact on sleep behavior to understand the effects of sleep modulation on neurodegeneration. To that end, we analyzed whether expressing Tau in the photoreceptors using *GMR-GAL4* caused sleep disturbances by monitoring their sleep behavior **(Figure 1E)**. When comparing Tau^WT^ or Tau^R406W^ to controls (GFP), there was no significant difference in either the amount of average sleep per 24 hours (**Figure 1F**), the number of sleep bouts (**Figure 1G**), or the average length of each sleep bout **(Figure 1H)**. Detailed analyses of sleep divided in night and day showed that expression of Tau in the photoreceptors had minimum effects in their innate sleep behavior **(Figure S2A-C)** and no significant change in number of sleep bouts **(Figure S2C)** or bout length **(Figure S2D)**. Together, this data suggests that expression of Tau in the photoreceptors causes minimum effect on sleep, making it a feasible model for us to examine the physiological impact of sleep modulation.

To establish a sleep modulation paradigm that induces significant changes in sleep time, we incorporated both sleep deprivation and sleep induction protocols. Sleep disruption was achieved using a mechanical “deprivator” that rotates 90° for a determined time[32]. During mechanical deprivation, we can observe significant reduction of sleep, followed by an earlier sleep onset and increased daytime sleep. These changes in sleep profile are due to the sleep homeostat attempting to reduce the amount of sleep need accumulated during a deprivation event. Sleep induction was achieved by gaboxadol feeding following previously published methods [17]. Flies were subjected to either mechanical sleep deprivation for two nights, 9 hours each night (Hours 111-120 and Hours 135-140) or gaboxadol feeding at 0.1mg/ml concentration for 4 days (Hours 48-156) **(Figure 2A)**. Sleep studies showed significant sleep behavior alterations after sleep deprivation or sleep induction in all groups **(Figure 2B)**. On average, sleep deprivation resulted in a reduction of total nighttime sleep by 160 min, 160 min and 95 min for GFP, Tau^WT^ and Tau^R406W^, respectively; while sleep induction resulted in an increase of nighttime sleep of 174 min, 185 min, 145 min, respectively. Daytime activity showed similar changes by sleep deprivation or induction **(Figure 2C)**. Detailed sleep bout analyses showed that the number of sleep bouts was decreased by sleep deprivation, while increased by sleep induction **(Figure 2D)**, without significant changes in sleep bout length **(Figure 2E)**. Taken together, these results showed successful sleep modulation after mechanical deprivation or sleep induction for all groups and demonstrated the feasibility of the optimized sleep modulation paradigm.

**Figure 2:**
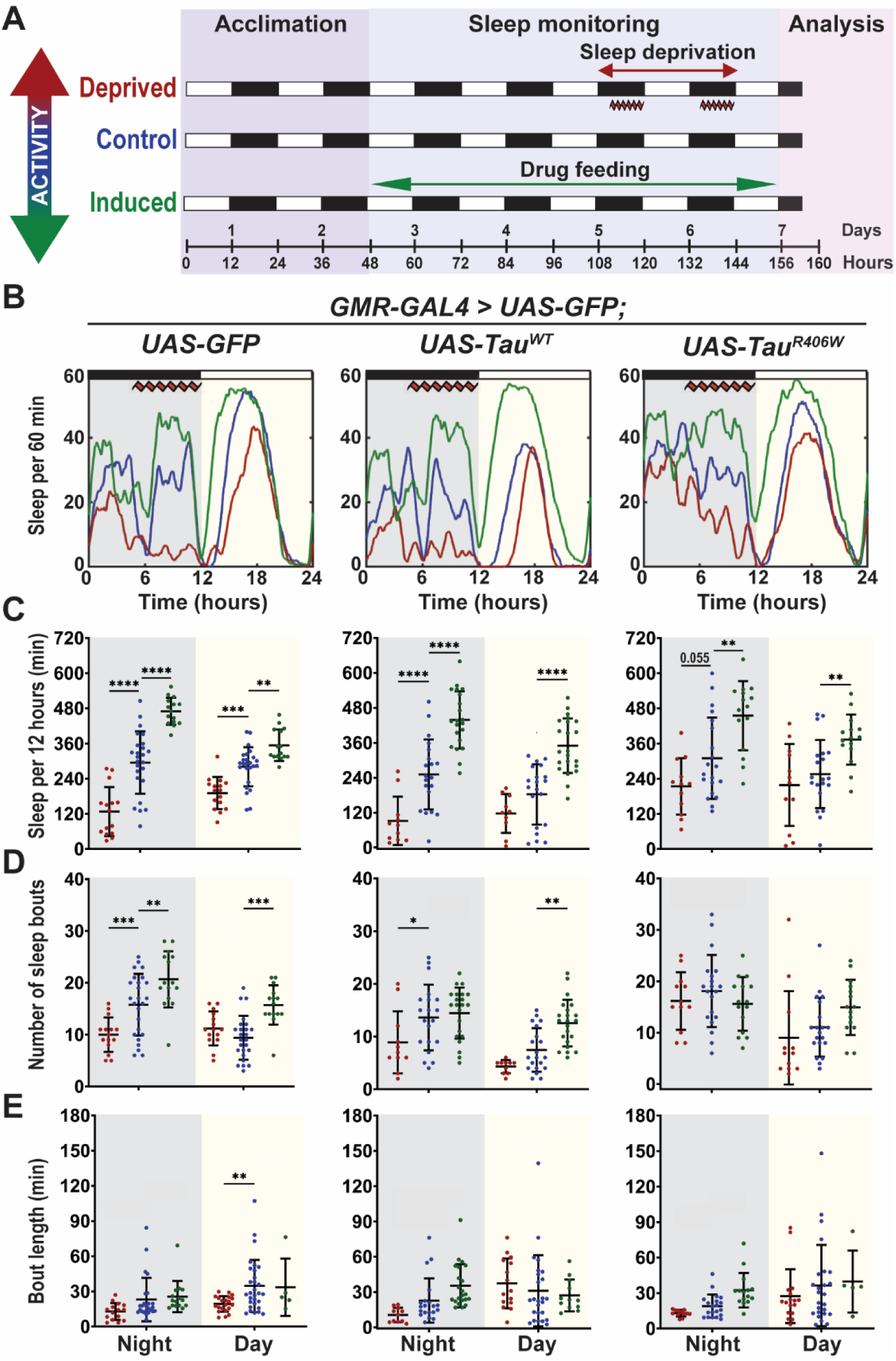
Established sleep paradigm successfully modulates total sleep time. **(A)** Sleep modulation paradigm. 2 DAE flies were placed in single glass tubes with food and their locomotor activity was monitored for 7 days. Deprived group was mechanically deprived for 9 hours on the last two nights of experiment (Hours 111-120 and Hours 135-144). Induced group was placed on food containing 0.1mg/mL gaboxadol for 4 days (Hours 48-144). Cellular and molecular analyses were performed at 160 hours. **(B)** Flies were expressing either *UAS-CD8-GFP, UAS-hTau^WT^ or UAS-hTau^R406W^* in the photoreceptors using *GMR-GAL4* driver. Sleep per 60 minutes traces showing control (blue), deprived (red), induced (green) groups. **(C)** Total sleep time per 12 hours during night (gray box) and day (yellow box) for Hours 132-156 is quantified. **(D)** Number of sleep bouts per 12 hours. **(E)** Sleep bout length in minutes per 12 hours. Data as mean ± SD. n = 10-25, One-way ANOVA, *\*P<0.05, **p<0.01, ***p<0.001****p<0.0001*.

It is important to note that all experiments were started using adult flies of 2 DAE (days after eclosion) and cellular and molecular analyses were performed 4 hours after the conclusion of sleep modulation (160 hours) on 8 DAE (**Figure 2A**). This paradigm was designed to examine the early effects of sleep modulation before the onset of severe neurodegeneration in Tauopathy models [28]. Collectively, combining pan-neuronal or cell type specific expression models with successful sleep modulation allows the mechanistic dissection of the impact of sleep modulation on neurodegeneration in *Drosophila* models of Tauopathy.

### Sleep modulation alters the progression of Tau-induced synaptic degeneration

Synaptic loss is one of the early events of neurodegeneration and main lesions presented in human AD patients [33]. We have previously reported that our Tau expression models recapitulate the phenotype of Tau induced synaptic degeneration [27]. To evaluate how sleep modulation affects synaptic integrity and function we expressed Tau^WT^ or Tau^R406W^ in the *Drosophila* visual system to take advantage of the highly organized parallel axons of the compound eye: the R1-R6 photoreceptors axons cross the lamina cortex and make synaptic connections at the lamina neuropil, while R7-R8 extended their axons beyond the lamina and project to the medullar neuropil [34, 35]. We examined how sleep modulation affects synaptic integrity by measuring Bruchpilot (BRP), an active zone associated cytomatrix protein using confocal imaging **(Figure 3A)** and synaptic function by using electroretinogram recordings (ERGs) **(Figure 3B)**.

**Figure 3:**
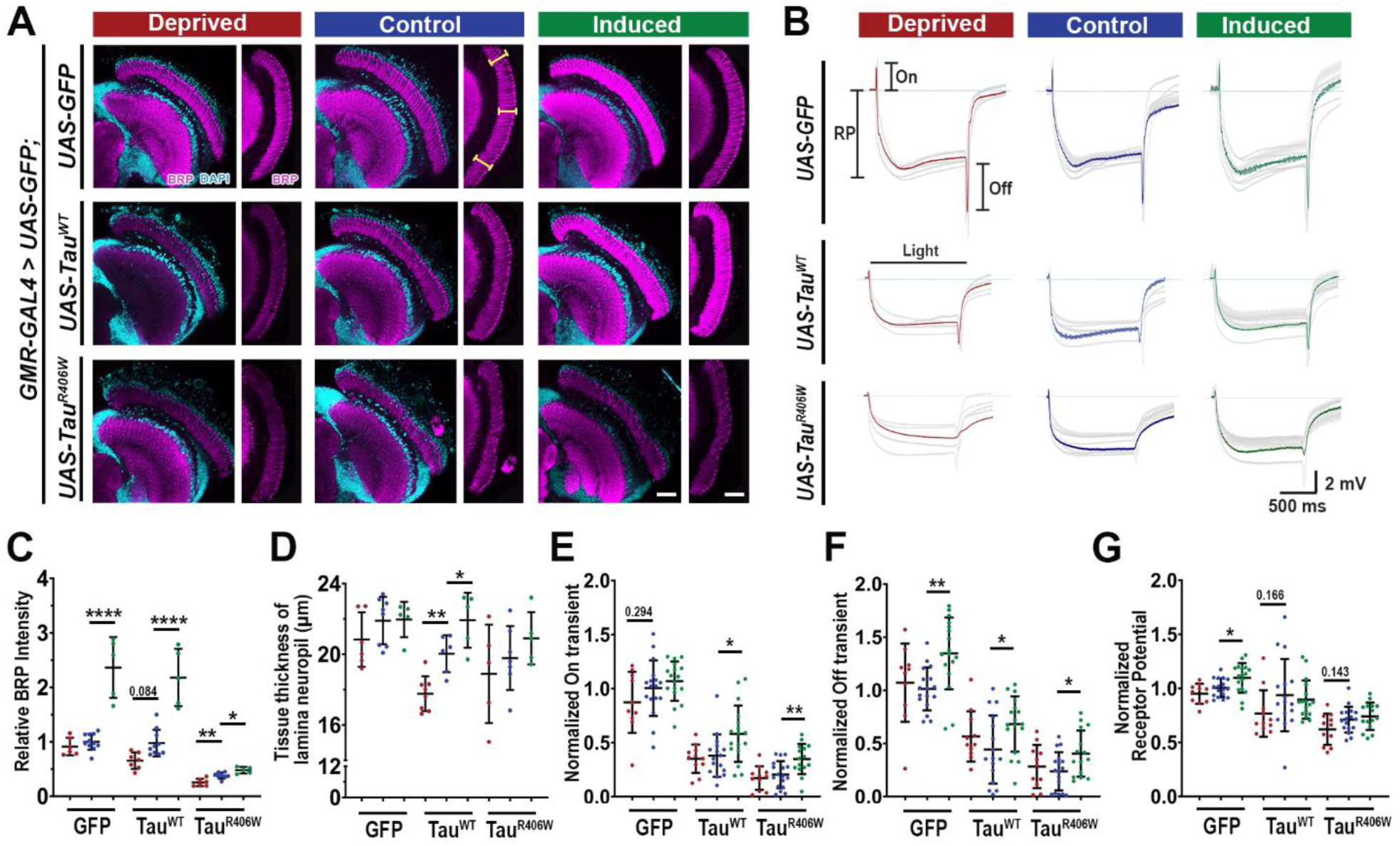
Sleep modulation alters the progression of tau-induced synaptic degeneration. **(A)** Lamina structures at 8 DAE probed for BRP (magenta) and stained with DAPI (cyan). Yellow dashed line highlights tissue thickness of lamina neuropil. Scale bar 30µm. Cellular and functional analyses were performed at 160 hours. **(B)** Electroretinogram recordings showing average response of 10 flies (gray). Average is shown in red, blue, and green respectively. RP: resting potential; On: on-transient; Off: off-transient. **(C)** Quantification of BRP intensity in the optic lobe normalized to GFP control. Brown-Forsythe and Welch ANOVA with Dunnett’s multiple comparison correction **(D)** Quantification of tissue thickness of lamina neuropil (yellow bars shown on panel A **(E-G)** Quantification of transients normalized to the GFP control. On transient **(E)**, Off transient **(F)** and Receptor potential **(G)** amplitude. For dissections n = 5-11 and for ERGs n≥10 light pulses recorded from 10-12 flies for each group. Data are presented as mean ± SD. One-way ANOVA, \**p<0.05*, \*\**p<0.01*, \*\*\**p<0.001*, \*\*\*\**p<0.0001*.

Measurement of total BRP intensity in the normal control (GFP) group showed no change after sleep deprivation but a ∼56% increase after sleep induction, suggesting a physiological effect of sleep induction on active zone BRP homeostasis. In Tauopathy groups (Tau^WT^ and Tau^R406W^), BRP level was reduced by 32% after sleep deprivation, and increased by 46-55% after sleep induction **(Figure 3C)**. These results suggest sleep deprivation exacerbated Tau-induced loss of active zone structures in the presynaptic terminals, while sleep induction significantly improved the localization of endogenous BRP at the synapses.

Consistent with BRP levels, the synaptic architectural integrity as indicated by tissue thickness of the lamina neuropil was also altered by sleep modulation. While sleep modulation had no significant effect on tissue thickness in normal control (GFP) group, sleep deprivation caused an ∼11.5% reduction, and sleep induction an ∼9.5% increase in tissue thickness of Tau^WT^. In Tau^R406W^, sleep deprivation caused a ∼4.6% reduction, while sleep induction showed a ∼5.7% increase in tissue thickness, on average **(Figure 3D)**. These results suggest that sleep deprivation exacerbated synaptic loss and sleep induction provided significant improvement of synaptic structures.

To examine the functional consequence of sleep modulation, we carried out electroretinogram (ERG) study of synaptic function. These electrical responses caused by light stimulation provide information about phototransduction (receptor potential, RP) as well as pre- and post-synaptic responses of the laminar monopolar cells (On and Off transients) [36–38]. Consistent with previous reports, Tauopathy flies showed reduced On- and Off-transients and receptor potential under control conditions, indicating synaptic degeneration [39]. Notably, these defects were partially ameliorated by sleep induction, which showed significant increases in the amplitude of On- and Off-transients, in all groups **(Figure 3E-3F).** On the other hand, sleep deprivation showed no significant ERG changes. The minimal change caused by sleep deprivation might indicate that a stronger or more prolonged deprivation effect is required to observe substantial functional degeneration at this early stage of disease. The improvements in On- and Off-transients after sleep induction suggests improved synaptic function, which is consistent with the overall improvement in synaptic morphology that was previously described. Similarly, deprivation did not result in RP amplitude deterioration, but sleep induction showed slight RP amplitude increase in normal control GFP flies **(Figure 3G)**, indicating that sleep induction has a general effect on all synapses.

Taken together, our morphological and functional study of the synapse demonstrates that sleep modulation is altering the progression of tau-induced synaptic degeneration. Specifically, sleep deprivation accelerates synaptic degeneration as shown by exacerbated loss of active zone proteins and overall synaptic structural integrity, while sleep induction improves overall synaptic physiology and structural integrity.

### Sleep modulation affects hyperphosphorylated Tau (pTau) accumulation in neurons

To dissect the molecular underpinnings of the effect of sleep modulation on Tau-induced synaptic degeneration, we set out to examine the biochemical changes of Tau. Expression of Tau^R406W^ results in Tau hyperphosphorylation and synaptic aggregation of hyperphosphorylated pTau [28]. We have optimized a high-resolution imaging approach that allows the characterization of the properties of pTau and the dynamic interaction with cellular organelles [28, 40]. Using pTau specific antibody AT8 (for Ser202, Thr205) that is specific for Tau pared helical fragments (PHF-tau), we quantitatively analyzed overall optic lobe pTau intensity and individual size distribution and intensity of pTau clusters to examine changes in the aggregational state of Tau caused by sleep modulation **(Figure 4A)**. The total intensity of pTau showed a remarkable alteration under sleep modulation with a significant increase after sleep deprivation and a modest reduction after sleep induction in Tau^R406W^ **(Figure 4B)**. When comparing individual pTau clusters a significant increase in intensity was observed after sleep deprivation for Tau^WT^ and a slight increase in Tau^R406W^ **(Figure 4C)**. The size analysis of pTau cluster revealed an interesting effect of sleep modulation, where sleep deprivation significantly increased medium and large clusters in Tau^R406W^ and Tau^WT^, respectively, while sleep induction decreased medium and large sized clusters in Tau^WT^ **(Figure 4D)**. To assess the filamentous accumulation of pTau in axons, we analyzed the circularity property of the pTau clusters and established a filamentous index, where the filamentous index is defined as the inverse of circularity using the formula: 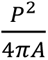, where A = Area and P = perimeter. The scale of filamentous index goes from 1, a perfect circle, to infinity, and the larger the number the more filamentous the pTau cluster. Interestingly, sleep deprivation increased the pool of high filamentous clusters (index >10), and sleep induction significantly decreased the mean filamentous index of pTau clusters in Tau^WT^ **(Figure 4E)**.

**Figure 4:**
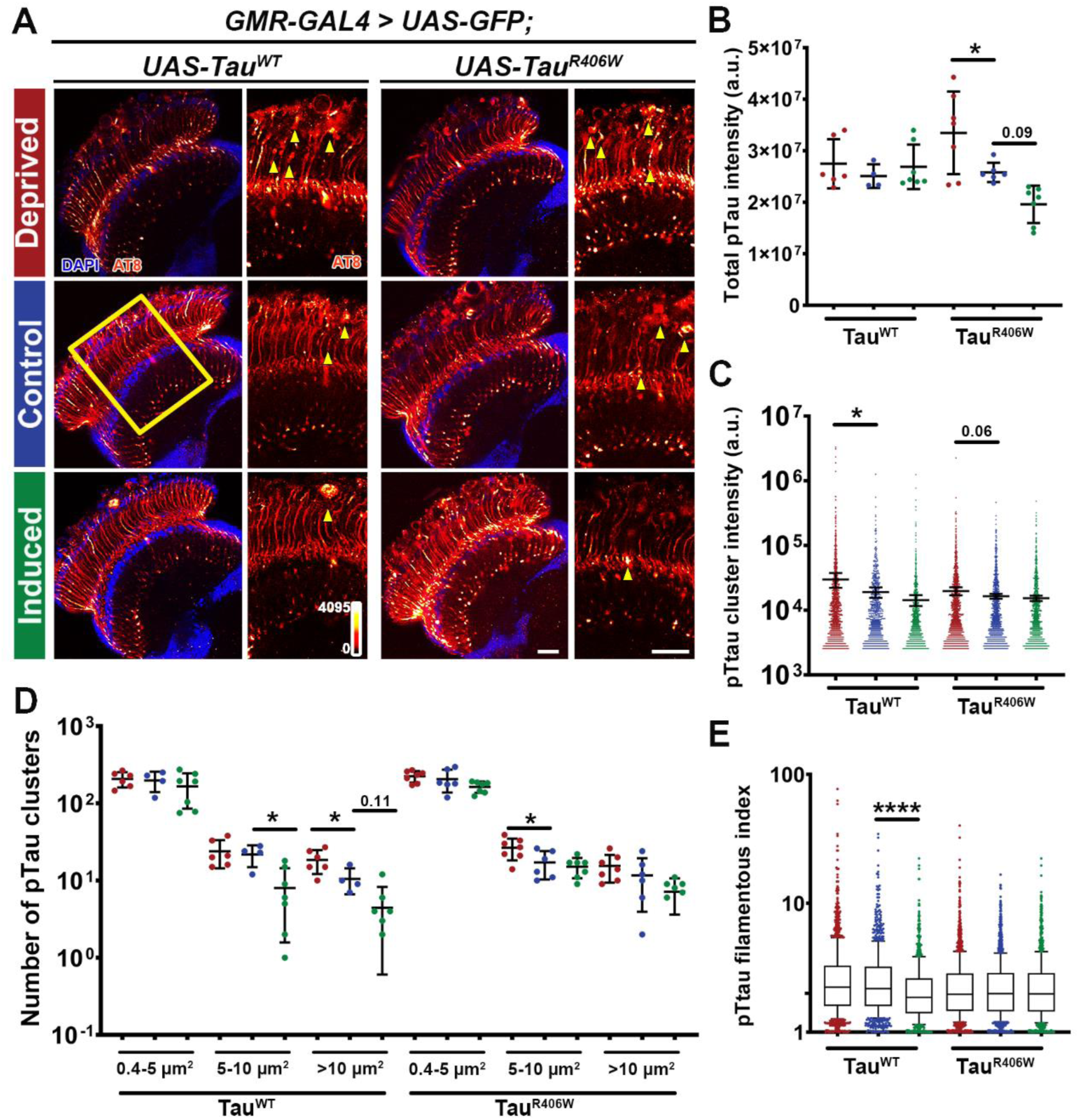
Sleep modulation alters hyperphosphorylated tau accumulation in neurons. **(A)** *Drosophila* optic lobe was stained with hyperphosphorylated tau antibody AT8 (Ser^202^, Thr^205^) (heatmap 0-4095). Yellow box represents zoomed in region of interest used for quantification. Scale bar 30µm. Yellow arrowheads represent AT8 clusters quantified in panels C and D. **(B)** Quantification of total AT8 intensity of optic lobe. **(C)** Quantification of intensity of AT8 clusters. **(D)** Quantification of number of AT8 clusters divided in small (0.4<5 µm^2^), medium (5-10 µm^2^), and large (>10 µm^2^) clusters. **(E)** Quantification of AT8 filamentous accumulation in axons. Data as mean ± SD. n = 4-7, One-way ANOVA, *\*p<0.05, ****p<0.0001*.

Taken together, these results show that when compared to controls, Tauopathy flies subjected to sleep deprivation have a significantly higher accumulation of pTau deposition in the brain. In contrast, sleep induction reduced the accumulation of hyperphosphorylated pTau specifically in medium and large clusters and dampened filamentous pTau formation in axons.

### Sleep modulation alters Tau aggregation and clearance

To uncover the mechanism underlying the reduction of pTau accumulation and the protective effects of sleep induction, we assessed the impact of sleep induction on Tau protein levels. To enable biochemistry analysis, we expressed Tau pan-neuronally using *elav-GAL4*. We first subjected the flies to sleep modulation and analyzed their sleep profiles **(Figure 5A)**. Quantification of sleep per 12 hours showed significant sleep reduction after sleep deprivation for normal control GFP group, but showed no significant deprivation for Tau expressing groups. This is likely due to the already significant sleep reduction and fragmentation due to Tau expression. In contrast, all groups showed significant sleep increase after gaboxadol feeding **(Figure 5B)**, allowing detailed biochemical analysis of the potential neuroprotective effects of sleep induction. Western blot analysis of Tau expression levels **(Figure 5C)** showed no changes in protein levels in the deprived group but a significant decrease of total Tau (5A6) **(Figure 5D)** and hyperphosphorylated pTau, using S262 probe **(Figure 5E)** after sleep induction, consistent with behavioral data. The significant reductions in both total Tau and pTau levels through sleep induction suggest Tau protein clearance as a key mechanism underlying the beneficial effects of sleep increase.

**Figure 5:**
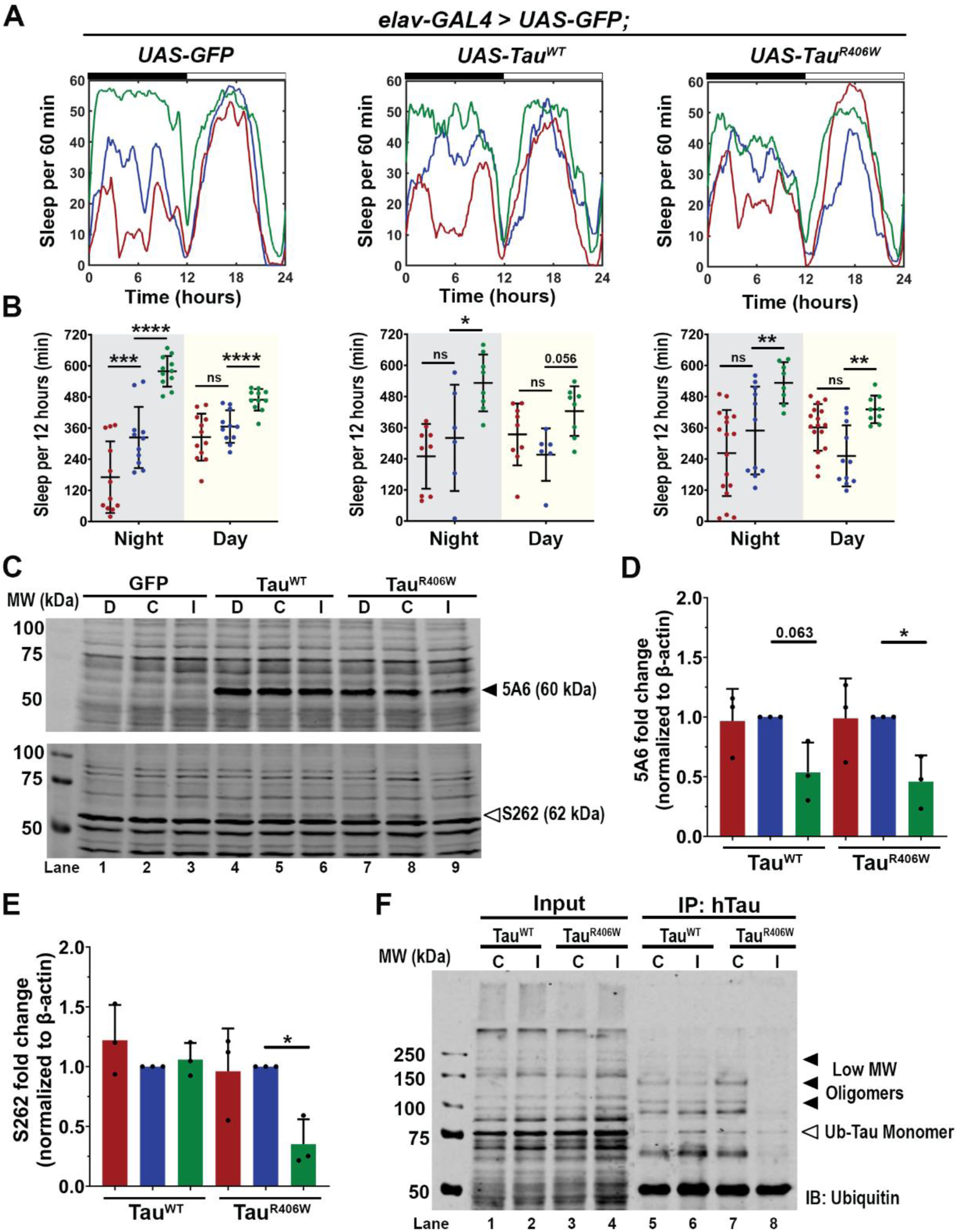
Sleep induction promotes clearance of ubiquitinated tau. **(A)** Sleep traces for 8 DAE flies expressing *UAS-GFP*, *UAS-hTau^WT^*or *UAS-hTau^R406W^* pan-neuronally using *elav-GAL4.* Sleep per 60 minutes traces showing control (blue), deprived (red), induced (green) groups. **(B)** Total sleep time per 12 hours during night (gray box) and day (yellow box) for Hours 132-156 is quantified. Data as mean ± SD. n = 6-17 flies, One-way ANOVA, *\*p<0.05*, \*\**p<0.01*, ***p<0.001, *\*\*\*\*p<0.0001*. **(C)** Western blot probed for total tau (5A6) (black arrowhead) and hyperphosphorylated tau (S262) (white arrowhead). **(D)** Quantification of 5A6 fold change normalized to β-actin. **(E)** Quantification of S262 fold change normalized to β-actin. . n = 3 biological replicates with 10 fly heads per group **(F)** Total hTau was immunoprecipitated from whole brain lysates of 8 DAE flies expressing either *Tau^WT^* or *Tau^R406W^* under control (c) or induction (I) and probed for ubiquitin. Blot shows low molecular weight (LMW) oligomers (black arrowheads) that can be mono- or polyubiquitinated and ubiquitinated monomers (white arrowhead). n = 30 fly heads per group Data as mean ± SD, One-way ANOVA, *\*p<0.05*, \*\**p<0.01*, *\*\*\*\*p<0.0001*.

Misfolded Tau proteins are usually degraded through multiple pathways such as the autophagy-lysosomal system and the ubiquitin-proteosome system [5]. A common early step in these degradation pathways is the ubiquitination of Tau to mark the protein for degradation. Our previous study has shown the different oligomeric states of ubiquitinated Tau and identified the strong toxicity associated with low molecular weight Tau oligomers [27]. We next examined the effects of sleep induction on the oligomerization of Tau species using biochemical approaches. Tau was immunoprecipitated by human Tau-specific antibody from lysates of Tau^WT^ or Tau^R406W^ expressing brains and probed from ubiquitin. Western blot analysis identified ubiquitinated Tau oligomeric species and found a reduction in low molecular weight toxic oligomers after sleep induction, when compared to control **(Figure 5F).** These results suggest that sleep induction alters Tau aggregation and promotes Tau clearance.

### Tau expression impairs autophagic flux

A large body of literature has demonstrated the role of autophagy in neurodegeneration and highlighted the disturbance of autophagic flux in AD and related disorders [41–43]. To assess how sleep induction impacts autophagic flux in Tauopathy models, we incorporated an in vivo autophagy dual-color reporter expressing Atg8 tandem tagged with pH sensitive GFP and mCherry at its amino terminal (*UAS-mCherry-GFP-Atg8a*) [44]. Since GFP signal is quenched under the acidic environment, combination of GFP and mCherry signal intensity will mark the stages of autophagy flux, where autophagosomes will be high GFP and high mCherry, while fusion with the lysosome will be low GFP and high mCherry **(Figure 6A)**.

**Figure 6:**
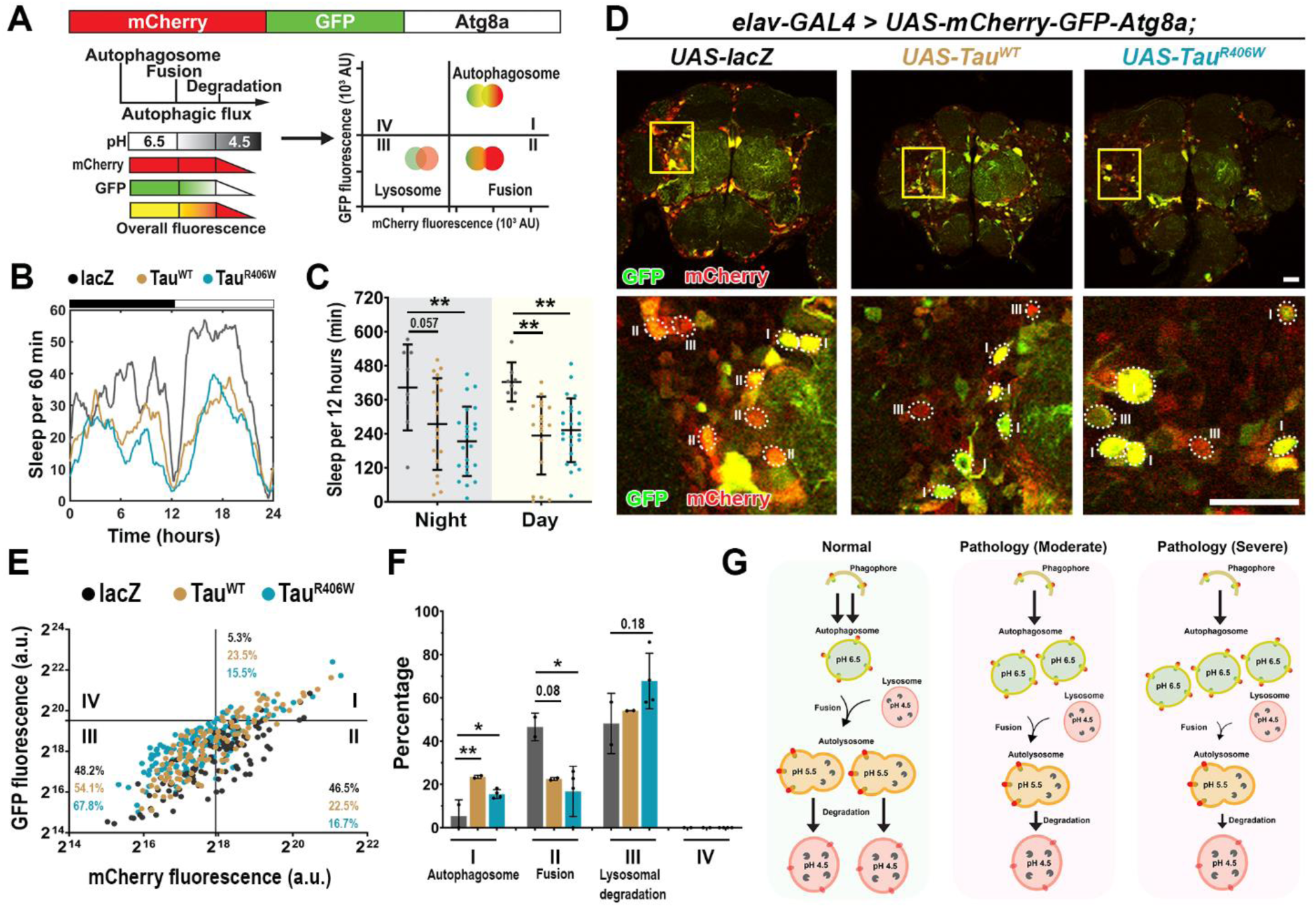
Pan-neuronal Tau expression displays impaired autophagic flux. **(A)** Model of fluorescent reporter with an mCherry (red fluorescent protein), GFP (green fluorescent protein) and Atg8a (autophagy fusion protein). GFP expression is quenched under acidic pH. Fusion requires low pH therefore, cells undergoing autophagy will display more puncta marked with mCherry-Atg8a (autophagosomes and autolysosomes) than with GFP-Atg8 (autophagosomes only). **(B)** Sleep profiles for 8 DAE flies with pan-neuronal expression using *elav-GAL4* of *UAS-mCherry-GFP-Atg8a* fluorescent reporter with *UAS-lacZ* (gray), *UAS-Tau^WT^* (gold) and *UAS-Tau^R406W^* (light blue). **(C)** Total sleep time per 12 hours during night (gray box) and day (yellow box) for Hours 132-156 is quantified. Data as mean ± SD. n = 8-23, One-way ANOVA, \*\**p<0.01*. **(D)** Confocal images of midbrain region showing GFP and mCherry fluorescence. Yellow box shows zoomed in cell body region. Dotted circles highlight Atg8a puncta. Scale bar 30µm. **(E)** mCherry-GFP-Atg8a puncta plotted by mean GFP intensity as a function of mean mCherry intensity. The plot was divided into four quadrants using gating thresholds of GFP = 750000 and mCherry = 250000. **(F)** Quantification of percentage of puncta in each quadrant; autophagosome stage (I), fusion (II) or lysosomal degradation (III). **(G)** Model showing that moderate and severe tauopathy impairs autophagic flux, causing increased autophagosome accumulation, decreased autophagosome -lysosome fusion and decreased autolysosomes degradation. Data as mean ± SD. n = 97-158 neurons from 3-4 fly brains, One-way ANOVA, *\*p<0.05*, \*\**p<0.01*.

We established autophagy reporter in Tauopathy models by co-expressing *UAS-mCherry-GFP-Atg8a* with either *UAS-lacZ* (control) *UAS-Tau^WT^* or *UAS-Tau^R406W^* pan-neuronally using *elav-Gal4.* We first analyzed the baseline sleep profiles to assess whether reporter expression caused changes in sleep behavior **(Figure 6B)**. Analysis of their activity behavior showed a significant reduction of nighttime and daytime sleep for Tau expressing groups when compared to lacZ **(Figure 6C)**. The activity profile is similar to the pan-neuronal Tauopathy models (**Figure 1A-1D**), suggesting that pan-neuronal Tau expression was the major driver of sleep disturbances, and the Atg8a reporter expression had minimum impact on sleep in Tauopathy models.

Using confocal imaging and ratiometric-based quantification [45] we analyzed the autophagic flux of Atg8 positive puncta at the neuronal cell body region of the midbrain as described previously [45] **(Figure 6D)**. The GFP and mCherry fluorescence intensity of all mCherry positive puncta was quantified and plotted in XY for the GFP fluorescence (Y) as a function of mCherry fluorescence (X) **(Figure 6E)**. Quadrants were divided based on intensity, where Quadrant I, high GFP-high mCherry marks autophagosome; Quadrant II, low GFP-high mCherry marks autolysosome; and Quadrant III, low GFP-low mCherry marks lysosomal degradation. The gating thresholds for quadrants were set based on the control group using the following principles: (1) The design of tandem reporter of pH-sensitive GFP and pH-insensitive mCherry defines the stoichiometry of GFP/mCherry as ≤ 1. Therefore, the content of Quadrant IV (GFP/mCherry ≥ 1) is 0. (2) The distribution of Atg8-positive particles reflects the distribution of autophagy-lysosome pathway related compartments in wild-type neurons [45, 56].

Compared to the lacZ control group, Tau^WT^ and Tau^R406W^ expressing neurons showed a significant increase in Quadrant I, autophagosome accumulation, and a significant reduction in Quadrant II, fusion of the autophagosome with the lysosome **(Figure 6F)**. These results reveal a shift in Atg8a puncta distribution in Tau pathology, and further suggest a potential block in autophagy flux at the step of fusion of autophagosome and lysosome **(Figure 6G)**. The finding of reduced fusion and a consequent increase in autophagosome accumulation and decreased fusion and degradation, would be consistent with Tau^WT^ or Tau^R406W^ expression-induced impairment of autophagic flux.

### Sleep improvement through promotion of autophagy

To examine the cellular impact of sleep modulation on autophagy flux, we first evaluated the control condition where lacZ and mCherry-GFP-Atg8 reporter were co-expressed. Sleep profiles confirmed that sleep modulation was successful **(Figure 7A)**, with significant sleep decrease after sleep deprivation and significant increased sleep after induction **(Figure 7B)**. Quantitative confocal imaging analysis of Atg8a puncta in the neuronal cell body region in the midbrain **(Figure 7C)**, showed significant shift in puncta distribution under sleep induction and minor changes under sleep deprivation **(Figure 7D-7E)**. Specifically, sleep induction caused a remarkable drop in Quadrant II and a concomitant increase in Quadrant III, suggesting a significant reduction in puncta in the fusion state, and a significant increase in puncta undergoing lysosomal degradation, **(Figure 7E)**. These results suggest that under normal (non-pathological) conditions, the effects of sleep deprivation on autophagy are minimal, while sleep induction appears to promote autophagic flux and may facilitate lysosomal degradation **(Figure 7F)**.

**Figure 7:**
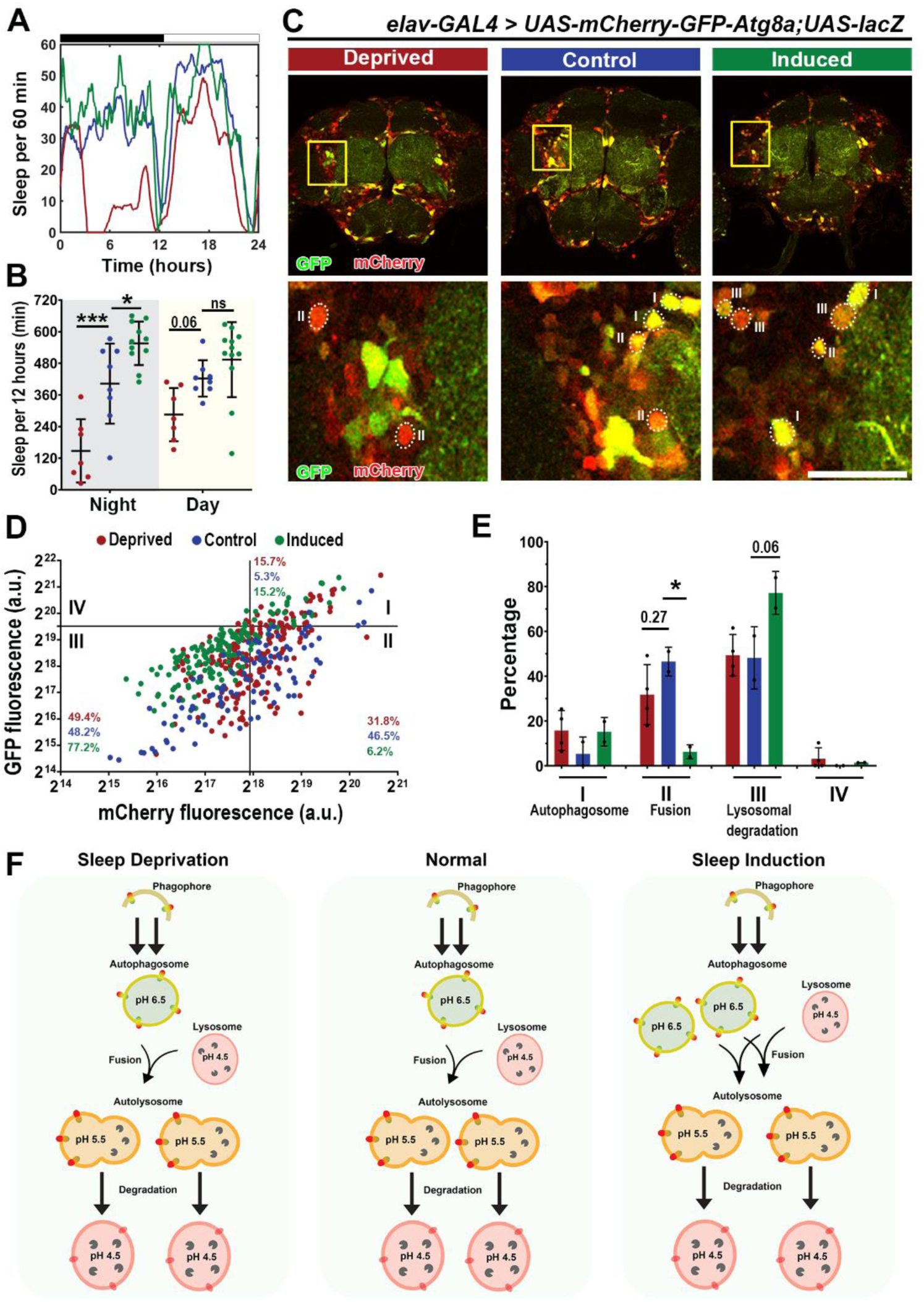
Sleep modulation promotes autophagic flux in lacZ. **(A)** Sleep traces for 8 DAE flies with pan-neuronal expression using *elav-GAL4* of *UAS-mCherry-GFP-Atg8a* fluorescent reporter with *UAS-lacZ* under control (blue), deprived (red) or induced (green) conditions. **(B)** Total sleep time per 12 hours during night (gray box) and day (yellow box) for Hours 132-156 is quantified. Data as mean ± SD. n = 8-12, One-way ANOVA, \**p<0.05*, \*\*\**p<0.001*. **(C)** Confocal images of midbrain region showing GFP and mCherry fluorescence. Yellow box shows zoomed in cell body region. Dotted circles highlight Atg8a puncta. Scale bar 30µm. **(D)** mCherry-GFP-Atg8a puncta plotted by mean GFP intensity as a function of mean mCherry intensity. The plot was divided into four quadrants using gating thresholds of GFP = 750000 and mCherry = 250000 for lacZ group. **(E)** Quantification of percentage of puncta in each quadrant for lacZ groups. **(F)** Model showing that under physiological conditions you have clearance of proteins through autophagy, while under sleep deprivation we do not observe significant changes in autophagic flux, but under sleep induction we observe increased flux, by increased fusion and degradation. Data are presented as mean ± SD. n = 97-212 neurons from 3-4 fly brains, One-way ANOVA, \**p<0.05*.

Next, we analyzed how sleep modulation affects autophagic flux in Tauopathy models. In Tau^WT^ expression (mild) model, sleep modulation was successful with sleep induction **(Figure 8A-8B)**. Imaging analysis of the Atg8a dual reporter expressed in the brain **(Figure 8C)** showed a significant increase in autophagosome puncta (Quadrant I) after sleep deprivation **(Figure 8D-8E)**. These results suggest that under moderate pathology sleep deprivation leads to increased autophagosome accumulation, while minimal changes were observed after sleep induction **(Figure 8F)**. In Tau^R406W^ expression (severe) model, sleep modulation was successful with sleep induction **(Figure 9A-9B),** similar to that in Tau^WT^ expression (mild) model (**Figure 8A**). However, analysis of autophagy flux using the Atg8a reporter revealed differences between the two pathological states **(Figure 9C)**. Specifically, no significant changes were observed after sleep deprivation, while after sleep induction, we observed an increase in autophagosome puncta **(Figure 9D-9E)**. These results suggest that under severe Tauopathy pathology, sleep induction leads to increased autophagy initiation, as shown by increased autophagosomes, which is expected to allow for more Tau to be degraded through autophagy **(Figure 9F)**.

**Figure 8:**
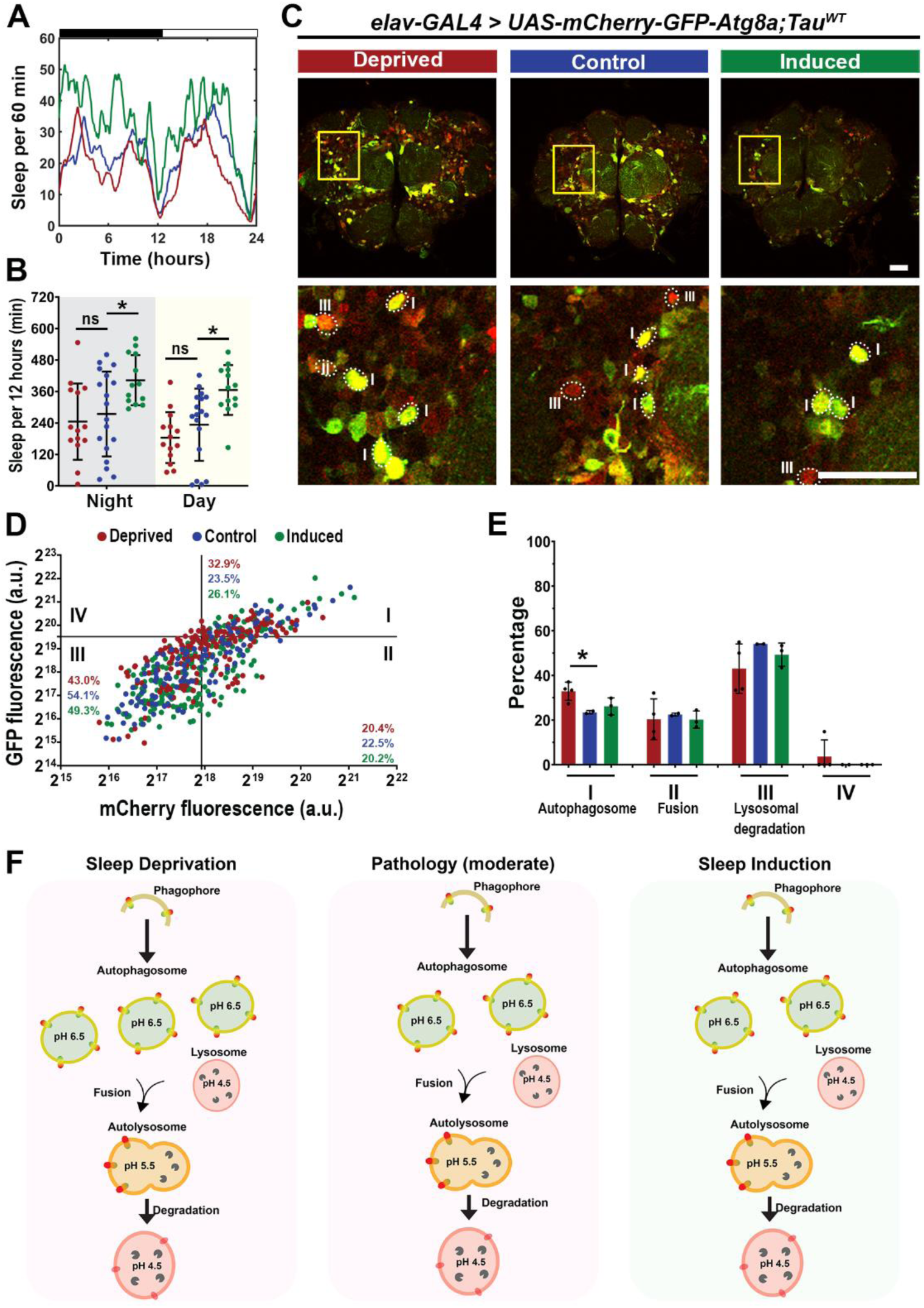
Sleep induction modulates autophagic flux in Tau^WT^ expression flies. **(A)** Sleep traces for 8 DAE flies with pan-neuronal expression using *elav-GAL4* of *UAS-mCherry-GFP-Atg8a* fluorescent reporter with *UAS-Tau^WT^*under control (blue), deprived (red) or induced (green) conditions. **(B)** Total sleep time per 12 hours during night (gray box) and day (yellow box)) for Hours 132-156 is quantified.. Data as mean ± SD. n = 13-18, One-way ANOVA, \**p<0.05*. **(C)** Confocal images of midbrain region showing GFP and mCherry fluorescence. Yellow box shows zoomed in cell body region. Dotted circles highlight Atg8a puncta. Scale bar 30µm. **(D)** mCherry-GFP-Atg8a puncta plotted by mean GFP intensity as a function of mean mCherry intensity. **(E**) Quantification of percentage of puncta in each quadrant. **(F)** Model showing moderate pathology impairs autophagic flux, increasing autophagosome accumulation and decreasing fusion of autolysosomes. Under sleep deprivation we observe exacerbated impairment, as shown by increased autophagosome accumulation. Data are presented as mean ± SD. n = 111-215 neurons from 3-4 fly brains, One-way ANOVA, \**p<0.05*.

**Figure 9:**
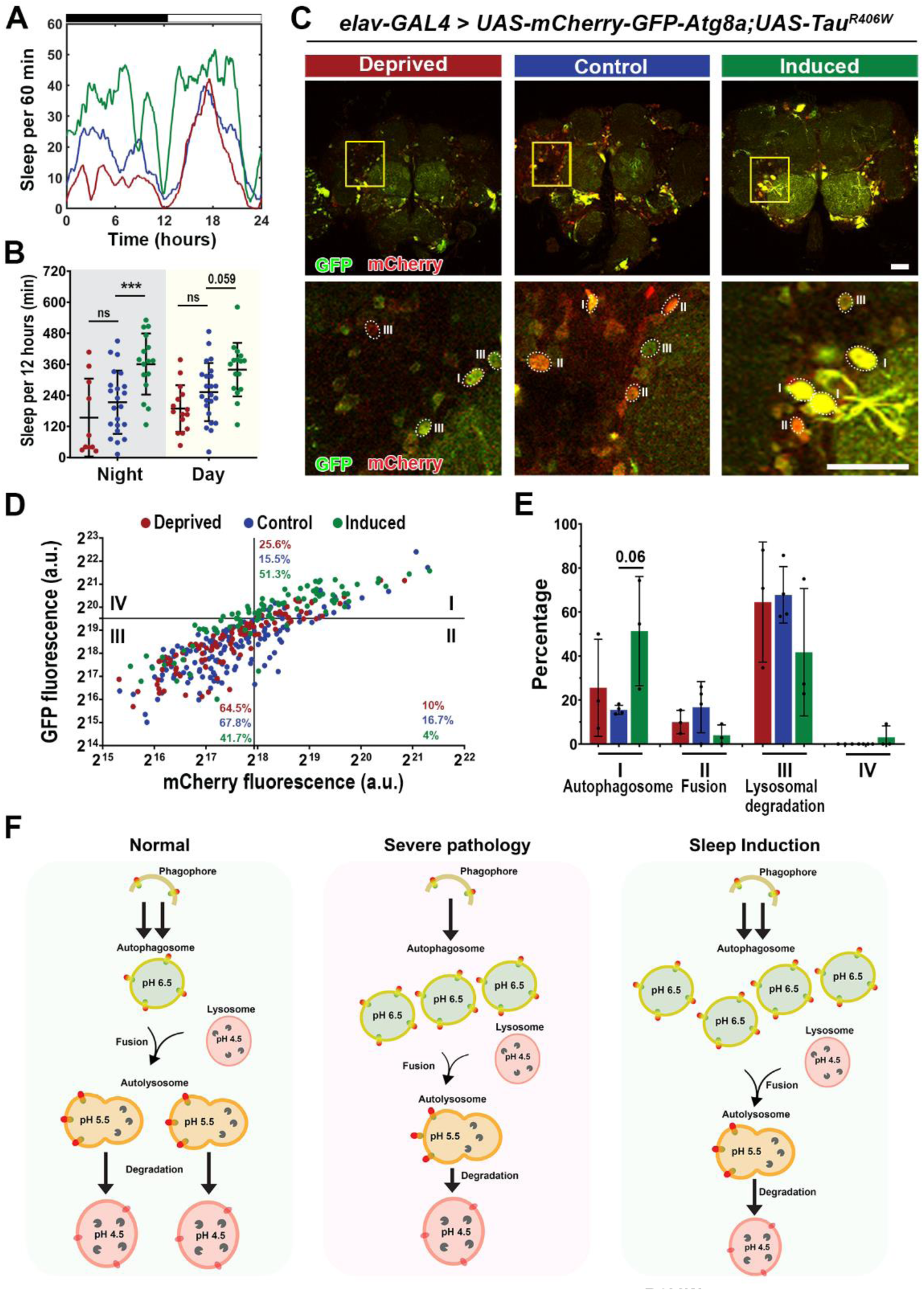
Sleep induction modulates autophagic flux in Tau^R406W^ expression flies. **(A)** Sleep traces for 8 DAE flies with pan-neuronal expression using *elav-GAL4* of *UAS-mCherry-GFP-Atg8a* fluorescent reporter with *UAS-Tau^R406W^*under control (blue), deprived (red) or induced (green) conditions. **(B)** Total sleep time per 12 hours during night (gray box) and day (yellow box) for Hours 132-156 is quantified. Data as mean ± SD. n = 10-23, One-way ANOVA, \*\*\**p<0.001*. **(C)** Confocal images of midbrain region showing GFP and mCherry fluorescence. Yellow box shows zoomed in cell body region. Dotted circles highlight Atg8a puncta. Scale bar 30µm. **(D)** mCherry-GFP-Atg8a puncta plotted by mean GFP intensity as a function of mean mCherry intensity. **(E**) Quantification of percentage of puncta in each quadrant. **(F)** Model showing that under severe pathology you have impaired autophagic flux, shown as increased autophagosome accumulation and decreased fusion of autolysosomes and with improvement in autophagic flux after sleep induction. Data are presented as mean + SD. n = 96-158 neurons from 3-4 fly brains, One-way ANOVA, *p<0.05.

## Discussion

In this study, we established sleep modulation paradigms to examine the effect of altered sleep on the progression of Tau-induced neurodegeneration in *Drosophila* models of Tauopathy. Using a multi-disciplinary approach of sleep behavior studies, genetic approaches, cellular and biochemical analysis, and physiological recordings, we discovered that sleep modulation alters Tau protein aggregation, clearance, and synaptic degeneration in vivo. We show that sleep deprivation enhanced Tau aggregational toxicity resulting in exacerbated synaptic degeneration. In contrast, sleep induction using gaboxadol led to reduced hyperphosphorylated Tau accumulation in neurons as a result of modulated autophagic flux and enhanced clearance of ubiquitinated Tau, suggesting altered protein processing and clearance that resulted in improved synaptic integrity and function. Specifically, pathological Tau expression impairs autophagic flux, accumulated hyperphosphorylated tau clusters and overall decreased synaptic integrity and function.

After sleep deprivation, there is increased autophagosome accumulation, increased medium and large sized pTau clusters that lead to exacerbated synaptic degeneration. In contrast, sleep induction exerts neuroprotective effects as a result of increased autophagosome initiation, decreased medium and large sized pTau clusters and improved synaptic integrity and function **(Figure 10)**.

**Figure 10:**
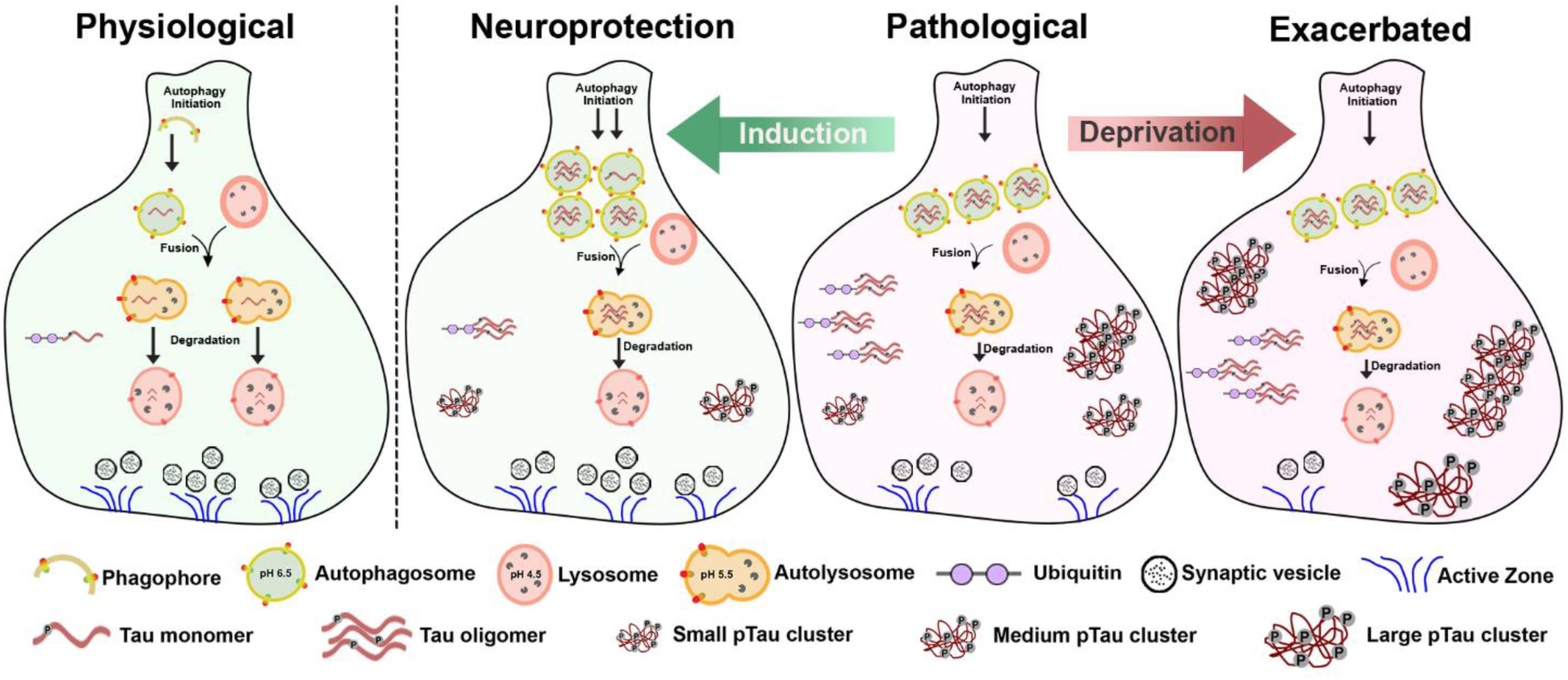
Sleep modulation alters tau aggregation, clearance, and synaptic degeneration in vivo in *Drosophila* models of Tauopathy. Schematic model of synapse showing clearance of proteins through autophagy under physiological conditions (Physiological). Pathological Tau expression results in impaired autophagic flux, accumulated hyperphosphorylated Tau clusters and overall decreased synaptic integrity and function (Pathological). Sleep deprivation leads to increased autophagosome accumulation and increased medium and large sized pTau clusters that result in exacerbated synaptic degeneration (Exacerbated). In contrast, sleep induction confers neuroprotective effects as a result of increased autophagosome initiation, decreased medium and large sized pTau clusters and improved synaptic integrity and function (Neuroprotection).

Sleep is essential for normal human function. It modulates synaptic plasticity; it is critical for memory consolidation, and it is essential for neurotoxic clearance of proteins from the brain [46]. Lack of sleep can disturb circadian physiology and lead to impaired memory, elevated oxidative stress, and increased risk of disease [1, 12, 47]. In Alzheimer’s Disease, altered sleep occurs early and lead to increased risk for developing cognitive impairment and AD progression.

Tau pathology is highly associated with synaptic loss and altered synaptic function and synapse loss is the main correlate with cognitive decline in AD [48, 49]. In this study, we observed that sleep deprivation resulted in increased hyperphosphorylated tau accumulation in axons. These results are consistent with increased evidence showing that sleep deprivation leads to neurotoxic protein accumulation such as Aβ and Tau [18, 31, 50, 51]. This increased accumulation of Tau caused accelerated progression of synaptic degeneration as observed by the decreased thickness of the lamina neuropil as well as decrease presence of active zones at synapses. Taken together, these results further confirm the direct interplay between sleep disturbances and protein homeostasis. Our data supports increasing evidence suggesting a positive feedback loop were impairment of one mechanism, exacerbates the other and together can accelerate AD progression.

Furthermore, sleep induction using gaboxadol, a GABA_A_ receptor agonist, showed substantial reduction of total Tau protein levels as well as hyperphosphorylated Tau. In addition, we observed a reduction of filamentous Tau accumulation in synapses. This reduction resulted in improved thickness of the lamina neuropil, as well as overall presence of active zones and improved synaptic function. It is important to note that the effect of gaboxadol has been shown to be mainly through increasing sleep, rather than GABA activation. Specifically, knockdown of GABA_A_ receptors prevented long-term memory consolidation following gaboxadol administration [52]. These results support the idea that sleep enhancing treatments can delay Tau induced neurodegeneration.

Moreover, we observed that pan-neuronal Tau expression leads to impaired autophagic flux, shown by increased autophagosome accumulation and decreased fusion and lysosomal degradation. This data supports research showing disturbed clearance of autophagic vacuoles in AD brain and APP mouse models [41] as well as increased autophagosome accumulation caused by MAPT overexpression in mice [42].

Previous work has highlighted the connection between sleep and autophagy specifically showing that under unperturbed autophagy, sleep increases clearance of autophagosomes and/or decreases production of autophagosomes. [17]. Our data using the autophagy reporter provides evidence that under pathological conditions were autophagy is already disturbed, the effects of sleep modulation might differ from unperturbed conditions. Sleep induction using gaboxadol showed increased autophagosomes, as well as led to decreased amounts of ubiquitinated low molecular weight Tau toxic oligomers. Together, this data suggests that sleep induction is promoting the formation of autophagosomes to push the flux forward and allow for more Tau degradation to occur.

With current treatments neurodegeneration persists despite interventional treatment. Some clinical trials are now exploring the potential effects of sleep aids such as suvorexant to decrease the rate of Aβ accumulation in the brain [53]. Many of the studies have focused on sleep interventions with already established AD dementia, where pathology is already too advanced, making treatments ineffective against the progression of disease [54]. Our data suggest that sleep can affect proteostasis and the clearance of misfolded proteins, consequently the progression of disease from early stages. Future work is required to identify biomarkers that detect neuronal susceptibility to changes in protein homeostasis. Therefore, given the abundance of sleep-enhancing pharmaceuticals on the market, determining the neuroprotective effects of sleep-inducing agents on protein homeostasis before the onset of clinical symptoms is critical for early intervention.

## Materials and Methods

### *Drosophila* stocks and genetics

Flies were reared on cornmeal-molasses-yeast medium at 22°C, 65% humidity, with 12 h light / 12 h dark cycles. The following strains were used in this study: UAS-Tau^WT^, UAS-Tau^R406W^ obtained from Dr. Mel Feany [27]. UAS-GFP-mCherry-Atg8a, GMR-GAL4, and elav-GAL4 were obtained from Bloomington *Drosophila* Stock Center.

### Locomotor activity

Male flies were placed in single glass tubes as previously described [55]. Locomotion was recorded using activity monitors (DAM2, Trikineticks Inc., MA). Monitors were placed in an environmental chamber maintained at 25°C and 70-80% relative humidity. In all experiments flies were first entrained in 12-h light/ 12-h dark conditions for 2 days. After entrainment, locomotion was measured with 20s binning for 4 days. Data from monitors were visualized and processed with custom-written Matlab (mathWorks Inc., MA) scripts.

### Sleep modulation

Mechanical deprivation was adapted from previously described Sleep Nullifying Apparatus (SNAP) methods [32]. Flies were subjected to two episodes of 9-hrs of deprivation. For sleep induction flies were fed either vehicle or 0.1mg/ml gaboxadol that was mixed with normal fly food [17].

### *Drosophila* CNS immunostaining, confocal imaging, and analysis

Fly brains were dissected in phosphate-buffered saline (PBS, pH 7.4) as previously described [45]. Samples were fixed in 4% formaldehyde for 10 min and washed in PBS containing 0.4% v/v Triton X-100 (PBTX). Samples were then incubated with primary antibodies diluted in 0.4% PBTX with 5% normal goat serum at 4°C overnight, followed by incubation with secondary antibodies diluted in 0.4% PBTX with 5% normal goat serum at 4°C overnight, and DAPI (1:300, Invitrogen, D1306) staining at room temperature for 10 min. The following primary antibodies were used: mouse anti-BRP (Developmental Studies Hybridoma Bank, NC82,1:250), AT8 human PHF-Tau (Thermo Scientific, MN1020, 1:250). The following secondary antibodies were used: Alexa Fluor 555-conjugated anti-mouse (Invitrogen, A21422, 1:300) and Alexa Fluor 647 anti-mouse secondary antibody (Invitrogen, A21235, 1:300). The samples were mounted on glass slides with VECTASHIELD Antifade Mounting Medium (Vector Laboratories). Slides were imaged using an Olympus IX81 confocal microscope with 40x or 60x oil immersion objective lens with a scan speed of 8.0 µs per pixel and spatial resolution of 1,024 x1,024 pixels. Images were processed using FluoView 10-ASW (Olympus). Quantification of number, area and intensity of fluorescently tagged proteins was carried out using ImageJ/Fiji (1.53q) using previously published methods [45].

For filamentous index the inverse of circularity formula was used: 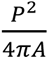, where A = Area and P = perimeter.

For quantification of autophagic flux using tandem reporter previously published methods were followed [45, 56]. The plot was divided into four quadrants using gating thresholds of GFP = 750000 and mCherry = 250000. The gating thresholds for quadrants were set based on the control group using the following principles: (1) The design of tandem reporter of pH-sensitive GFP and pH-insensitive mCherry defines the stoichiometry of GFP/mCherry as ≤ 1. Therefore, the content of Quadrant IV (GFP/mCherry ≥ 1) is 0. (2) The distribution of Atg8-positive particles reflects the distribution of autophagy-lysosome pathway related compartments in wild-type neurons [45, 56].

### Electroretinogram recordings and analysis

Previously described methods were followed [38, 57]. Flights were anesthetized with CO_2_ and immobilized on a glass slide. A recording electrode with 3M NaCl was placed on the surface of the left eye, and another reference electrode was inserted into the thorax. After 5 minutes of dark adaptation, flies were given 1-second light stimulation (Digitimer), with 5 seconds of dark between each stimulus a total of 10 traces were recorded per fly and analyzed by pCLAMP 10 Electrophysiology Data Acquisition & Analysis Software (ver.10.5).

### Immunoprecipitation and Western Blot analysis

Ten fly heads of each genotype were collected, and flash frozen before extraction. They were homogenized in radioimmunoprecipitation assay (RIPA) buffer (Sigma-Aldrich, St. Louis, MO,USA), protease inhibitor cocktail (Sigma) and a phosphatase inhibitor cocktail (Roche). For immunoprecipitation experiments, 30 8 DAE male fly heads were homogenized in lysis buffer, and incubated wit Protein-G beads conjugated with 10µg of antibody per sample overnight at 4°C the bead pellets were collected and washed four times with lysis buffer before eluting out the beads fraction with 4x Lamelli buffer, heating the samples at 95°C and centrifuging for 5 min to collect the bead-eluted fraction. Lysates were probed with anti-human tau (Developmental Studies Hybridoma Bank, 5A6, 1:1000). For western blotting, ten microliters of total protein lysate per sample (equivalent to 1 head) was loaded onto a polyacrylamide gel (PAGE) and resolved by SDS-PAGE and transferred onto nitrocellulose membrane. After blocking at room temperature for 1hr, the membrane was incubated with primary antibody at 4°C overnight, followed by secondary antibody for 1hr at room temperature. Imaging was performed on an Odyssey Infrared Imaging System (LI-COR Biosciences) and analyzed using Image studio (v4.0). Primary antibody dilutions were used as follows: ubiquitin (Cell signaling, 3936, 1:1000), 5A6 anti-human tau (Developmental Studies Hybridoma Bank, 5A6, 1:250), Phospho-Tau Ser^262^ (Thermo Scientific, 44-750G, 1:1,000), anti-β-actin (Sigma-Aldrich, A1978, 1:5,000). Secondary antibodies used were: IRDYE800CW anti-mouse (Rockland, 610-145-002, 1:10,000) and IRDYE700DX anti-rabbit (Rockland, 611-144-002, 1:10,000).

### Statistical analyses

Biological sample size (n) and p values are indicated in the corresponding figure legends. Either One-way ANOVA with Tukey post-hoc test or Brown-Forsythe and Welch ANOVA with Dunnett’s multiple comparison correction was applied to compare multiple groups. *p* < 0.05 was considered statistically significant. All statistical analyses were performed in GraphPad Prism software (version 10.0).

## Acknowledgments

We thank Zoraida Diaz-Perez for technical assistance.

## Funding

This work was supported in part by Florida Department of Health (FDOH 21A21) (RGZ), and the National Science Foundation (NSF# 2131037).

## Author contributions

Conceptualization: RGZ, SS

Methodology: NOV, AGL, YZ, SL, SS, RGZ

Investigation: NOV, AGL, YZ, SL, TC, SS, RGZ

Visualization: NOV

Supervision: RGZ, SS

Writing—original draft: NOV, RGZ

Writing—review & editing: NOV, AGL, TC, YZ, SL, SS, RGZ

## Competing interests

Authors declare that they have no competing interests.

## Data and materials availability

All data are available in the main text or the supplementary materials.

## Supplementary Materials

**Figure S1:**
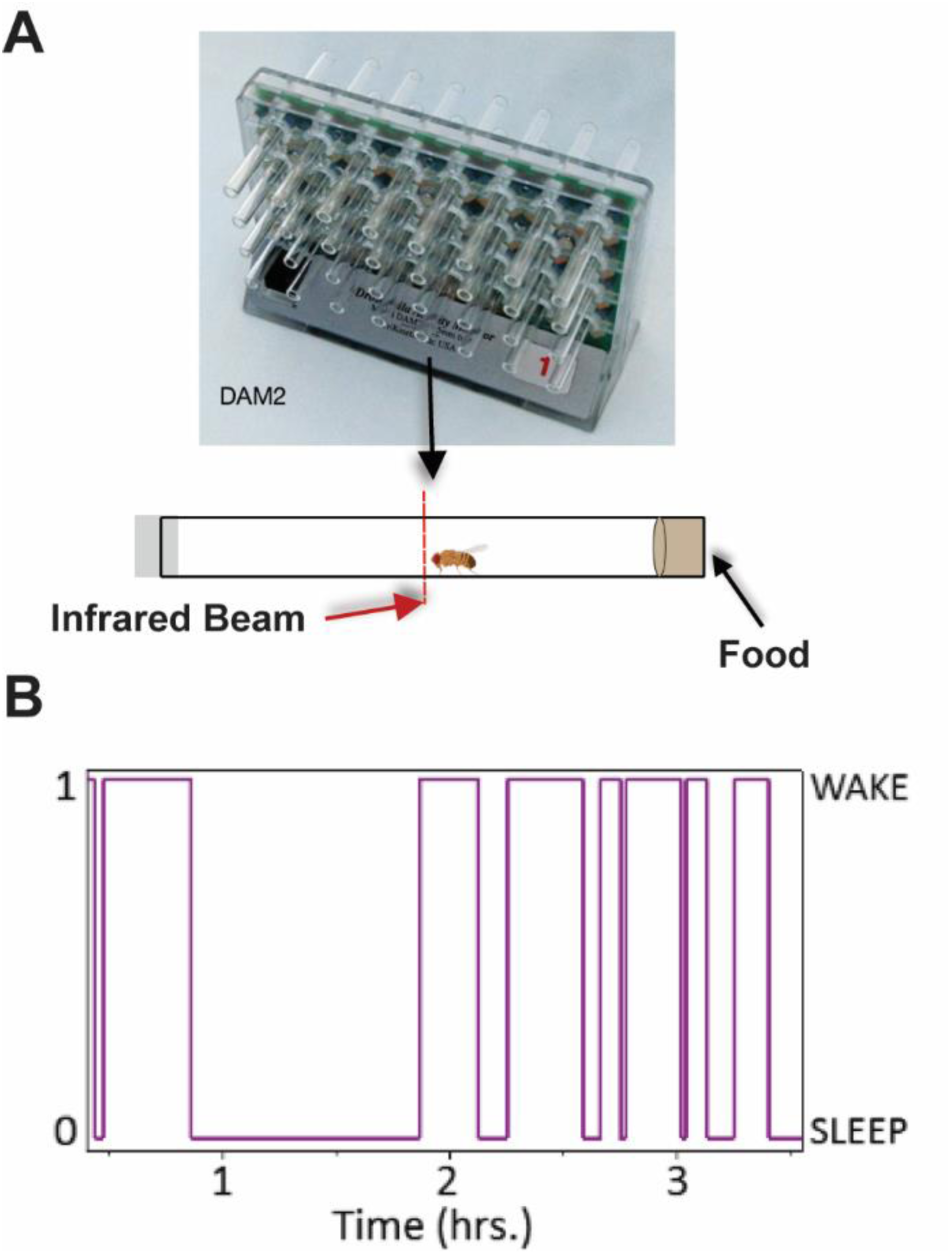
*Drosophila* activity monitoring. **(A)** Diagram showing flies placed in single glass tubes with food on one end and cotton stopper on the other end. Placed in a *Drosophila* activity monitoring (DAM) system with infra-red (IR) emitter detector pairs. **(B)** Example trace of an individual fly switching between sleep ‘0’ and wake ‘1’ states. Diagrams were adapted from Trikinetics and from “*Drosophila* male (lateral)”, by BioRender.com (2024). Retrieved from https://app.biorender.com/biorender-templates.

**Figure S2:**
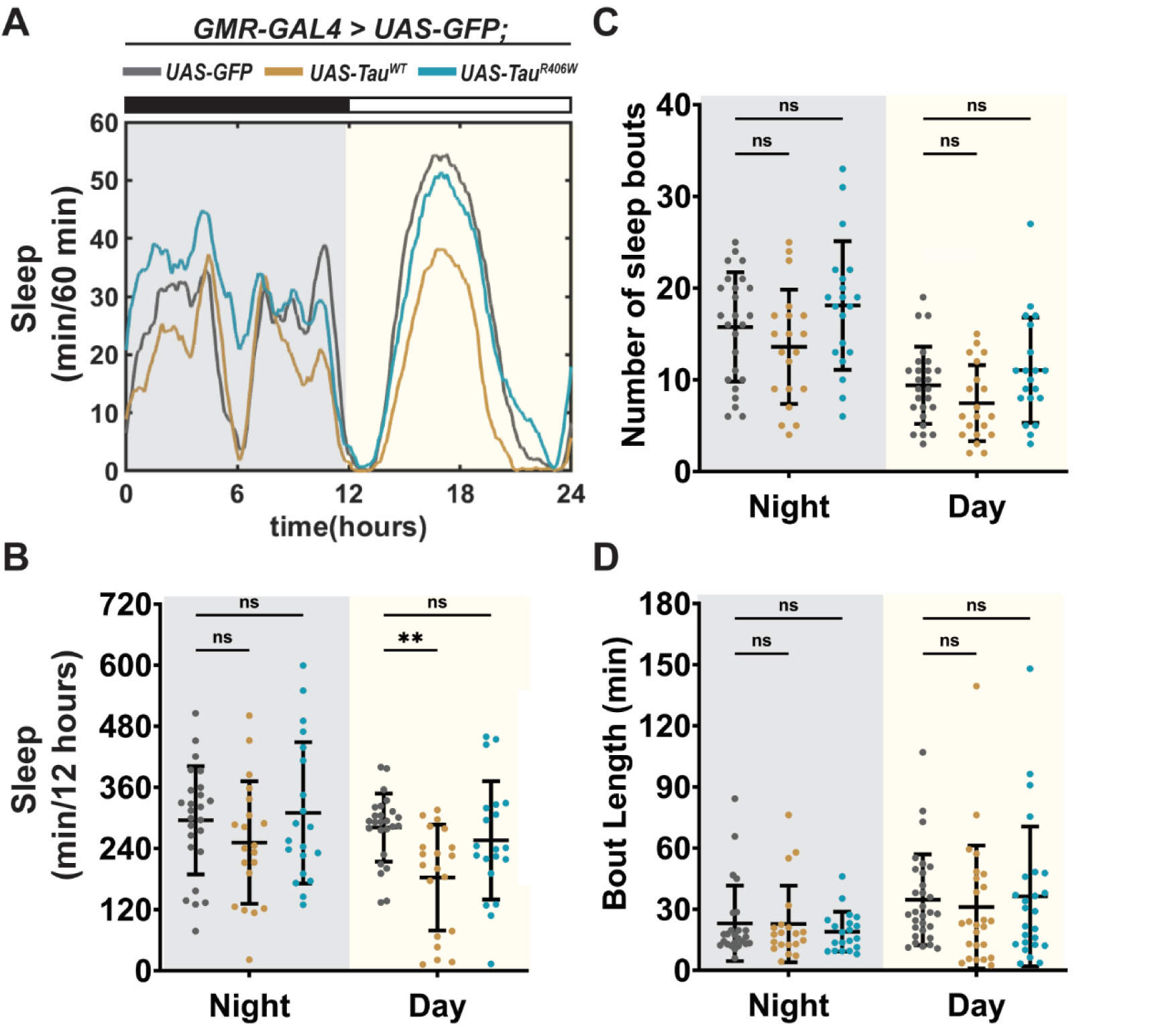
Expression of Tau in photoreceptors showed minimal effect on sleep behavior. Detailed analysis of Figure 1E, showing sleep quantification from Hours 132-156. Flies were expressing either *UAS-CD8-GFP* (gray), *UAS-hTau^WT^* (gold) or *UAS-hTau^R406W^*(turquoise) in the photoreceptors using *GMR-GAL4* driver. **(A)** Traces showing amount of sleep per 60 minutes in 24 hours. **(B)** Total sleep time per 12 hours during night (gray box) and day (yellow box). **(C)** Number of sleep bouts per 12 hours. **(D)** Sleep bout length in minutes per 12 hours. Data as mean ± SD. n = 20, One-way ANOVA, ** p< 0.01; ns, p>0.05.

## Notes

### Competing Interest Statement

The authors have declared no competing interest.

## References

1. Ju, Y.-E.S., B.P. Lucey, and D.M. Holtzman, Sleep and Alzheimer disease pathology—a bidirectional relationship. Nature Reviews Neurology, 2014. 10(2): p. 115–119.

2. Iranzo, A., Sleep in Neurodegenerative Diseases. Sleep Med Clin, 2016. 11(1): p. 1–18.

3. Alonso, A.D.C., I. Grundke-Iqbal, and K. Iqbal, Alzheimer’s disease hyperphosphorylated tau sequesters normal tau into tangles of filaments and disassembles microtubules. Nature Medicine, 1996. 2(7): p. 783–787.

4. Sperling, R.A., et al., Toward defining the preclinical stages of Alzheimer’s disease: Recommendations from the National Institute on Aging-Alzheimer’s Association workgroups on diagnostic guidelines for Alzheimer’s disease. Alzheimer’s & Dementia, 2011. 7(3): p. 280–292.

5. Jiang, S. and K. Bhaskar, Degradation and Transmission of Tau by Autophagic-Endolysosomal Networks and Potential Therapeutic Targets for Tauopathy. Frontiers in Molecular Neuroscience, 2020. 13.

6. Iqbal, K., et al., Tau in Alzheimer Disease and Related Tauopathies. Current Alzheimer Research, 2010. 7(8): p. 656–664.

7. Peter-Derex, L., et al., Sleep and Alzheimer’s disease. Sleep Med Rev, 2015. 19: p. 29–38.

8. Moran, M., et al., Sleep disturbance in mild to moderate Alzheimer’s disease. Sleep Med, 2005. 6(4): p. 347–52.

9. D’Rozario, A.L., et al., Objective measurement of sleep in mild cognitive impairment: A systematic review and meta-analysis. Sleep Med Rev, 2020. 52: p. 101308.

10. Mattis, J. and A. Sehgal, Circadian Rhythms, Sleep, and Disorders of Aging. Trends in Endocrinology & Metabolism, 2016. 27(4): p. 192–203.

11. Fifel, K. and A. Videnovic, Circadian and Sleep Dysfunctions in Neurodegenerative Disorders-An Update. Front Neurosci, 2020. 14: p. 627330.

12. Bishir, M., et al., Sleep Deprivation and Neurological Disorders. Biomed Res Int, 2020. 2020: p. 5764017.

13. Bubu, O.M., et al., Sleep, Cognitive impairment, and Alzheimer’s disease: A Systematic Review and Meta-Analysis. Sleep, 2017. 40(1).

14. Iliff, J.J., et al., A Paravascular Pathway Facilitates CSF Flow Through the Brain Parenchyma and the Clearance of Interstitial Solutes, Including Amyloid β. Science Translational Medicine, 2012. 4(147): p. 147ra111–147ra1.

15. Shokri-Kojori, E., et al., beta-Amyloid accumulation in the human brain after one night of sleep deprivation. Proc Natl Acad Sci U S A, 2018. 115(17): p. 4483–4488.

16. Spira, A.P., et al., Excessive daytime sleepiness and napping in cognitively normal adults: associations with subsequent amyloid deposition measured by PiB PET. Sleep, 2018. 41(12).

17. Bedont, J.L., et al., Short and long sleeping mutants reveal links between sleep and macroautophagy. eLife, 2021. 10.

18. Kang, J.-E., et al., Amyloid-β Dynamics Are Regulated by Orexin and the Sleep-Wake Cycle. Science, 2009. 326(5955): p. 1005–1007.

19. Sterniczuk, R., et al., Sleep disturbance is associated with incident dementia and mortality. Curr Alzheimer Res, 2013. 10(7): p. 767–75.

20. Dissel, S., Drosophila as a Model to Study the Relationship Between Sleep, Plasticity, and Memory. Front Physiol, 2020. 11: p. 533.

21. Fernandez-Funez, P., L. de Mena, and D.E. Rincon-Limas, Modeling the complex pathology of Alzheimer’s disease in Drosophila. Exp Neurol, 2015. 274(Pt A): p. 58–71.

22. Prussing, K., A. Voigt, and J.B. Schulz, Drosophila melanogaster as a model organism for Alzheimer’s disease. Mol Neurodegener, 2013. 8: p. 35.

23. Dubowy, C. and A. Sehgal, Circadian Rhythms and Sleep in Drosophila melanogaster. Genetics, 2017. 205(4): p. 1373–1397.

24. Hendricks, J.C., et al., Rest in Drosophila Is a Sleep-like State. Neuron, 2000. 25(1): p. 129–138.

25. Shaw, P.J., et al., Correlates of sleep and waking in Drosophila melanogaster. Science, 2000. 287(5459): p. 1834–7.

26. Pfeiffenberger, C., et al., Locomotor activity level monitoring using the Drosophila Activity Monitoring (DAM) System. Cold Spring Harb Protoc, 2010. 2010(11): p. pdb prot5518.

27. Ali, Y.O., K. Ruan, and R.G. Zhai, NMNAT suppresses Tau-induced neurodegeneration by promoting clearance of hyperphosphorylated Tau oligomers in a Drosophila model of tauopathy. Human Molecular Genetics, 2012. 21(2): p. 237–250.

28. Ma, X., et al., Nicotinamide mononucleotide adenylyltransferase uses its NAD+ substrate-binding site to chaperone phosphorylated Tau. eLife, 2020. 9.

29. Zhang, M.Y., B.C. Lear, and R. Allada, The microtubule-associated protein Tau suppresses the axonal distribution of PDF neuropeptide and mitochondria in circadian clock neurons. Hum Mol Genet, 2022. 31(7): p. 1141–1150.

30. Buhl, E., J.P. Higham, and J.J.L. Hodge, Alzheimer’s disease-associated tau alters Drosophila circadian activity, sleep and clock neuron electrophysiology. Neurobiol Dis, 2019. 130: p. 104507.

31. Tabuchi, M., et al., Sleep interacts with abeta to modulate intrinsic neuronal excitability. Curr Biol, 2015. 25(6): p. 702–712.

32. Melnattur, K., et al., The Sleep Nullifying Apparatus: A Highly Efficient Method of Sleep Depriving Drosophila. J Vis Exp, 2020(166).

33. Griffiths, J. and S.G.N. Grant, Synapse pathology in Alzheimer’s disease. Semin Cell Dev Biol, 2023. 139: p. 13–23.

34. Fischbach, K.-F. and A.P.M. Dittrich, The optic lobe of Drosophila melanogaster. I. A Golgi analysis of wild-type structure. Cell and Tissue Research, 1989. 258(3).

35. Katz, B. and B. Minke, Drosophila photoreceptors and signaling mechanisms. Frontiers in Cellular Neuroscience, 2009. 3(2).

36. Dolph, P., A. Nair, and P. Raghu, Electroretinogram recordings of Drosophila. Cold Spring Harb Protoc, 2011. 2011(1): p. pdb.prot5549.

37. Alawi, A.A. and W.L. Pak, On-Transient of Insect Electroretinogram: Its Cellular Origin. Science, 1971. 172(3987): p. 1055–1057.

38. Belusic, G., ERG in Drosophila, in Electroretinograms. 2011, InTech.

39. Fernandez-Funez, P. and R.R. Myers, Recent Contributions of the Drosophila Eye to Unraveling the Basis of Neurodegeneration, in Molecular Genetics of Axial Patterning, Growth and Disease in Drosophila Eye. 2020, Springer International Publishing. p. 293–309.

40. Brazill, J.M., et al., Quantitative Cell Biology of Neurodegeneration in <em>Drosophila</em> Through Unbiased Analysis of Fluorescently Tagged Proteins Using ImageJ. Journal of Visualized Experiments, 2018(138).

41. Manczak, M., et al., Hippocampal mutant APP and amyloid beta-induced cognitive decline, dendritic spine loss, defective autophagy, mitophagy and mitochondrial abnormalities in a mouse model of Alzheimer’s disease. Human Molecular Genetics, 2018. 27(8): p. 1332–1342.

42. Feng, Q., et al., MAPT/Tau accumulation represses autophagy flux by disrupting IST1-regulated ESCRT-III complex formation: a vicious cycle in Alzheimer neurodegeneration. Autophagy, 2020. 16(4): p. 641–658.

43. Nixon, R.A., et al., Extensive involvement of autophagy in Alzheimer disease: an immuno-electron microscopy study. J Neuropathol Exp Neurol, 2005. 64(2): p. 113–22.

44. Mauvezin, C., et al., Assays to monitor autophagy in Drosophila. Methods, 2014. 68(1): p. 134–139.

45. Brazill, J.M., et al., Quantitative Cell Biology of Neurodegeneration in Drosophila Through Unbiased Analysis of Fluorescently Tagged Proteins Using ImageJ. J Vis Exp, 2018(138).

46. Ly, S., A.I. Pack, and N. Naidoo, The neurobiological basis of sleep: Insights from Drosophila. Neurosci Biobehav Rev, 2018. 87: p. 67–86.

47. Hill, V.M., et al., A bidirectional relationship between sleep and oxidative stress in Drosophila. PLoS Biol, 2018. 16(7): p. e2005206.

48. Bejanin, A., et al., Tau pathology and neurodegeneration contribute to cognitive impairment in Alzheimer’s disease. Brain, 2017. 140(12): p. 3286–3300.

49. Coomans, E.M., et al., In vivo tau pathology is associated with synaptic loss and altered synaptic function. Alzheimer’s Research & Therapy, 2021. 13(1).

50. Spira, A.P., et al., Self-reported Sleep and β-Amyloid Deposition in Community-Dwelling Older Adults. JAMA Neurology, 2013.

51. Lucey, B.P., et al., Effect of sleep on overnight cerebrospinal fluid amyloid β kinetics. Annals of Neurology, 2018. 83(1): p. 197–204.

52. Dissel, S., et al., Sleep Restores Behavioral Plasticity to Drosophila Mutants. Current Biology, 2015. 25(10): p. 1270–1281.

53. Herring, W.J., et al., Polysomnographic assessment of suvorexant in patients with probable Alzheimer’s disease dementia and insomnia: a randomized trial. Alzheimer’s & Dementia, 2020. 16(3): p. 541–551.

54. Blackman, J., et al., Pharmacological and non-pharmacological interventions to enhance sleep in mild cognitive impairment and mild Alzheimer’s disease: A systematic review. J Sleep Res, 2021. 30(4): p. e13229.

55. Lazopulo, A. and S. Syed, A Computational Method to Quantify Fly Circadian Activity. Journal of Visualized Experiments, 2017(128).

56. Li, C., et al., Spermine synthase deficiency causes lysosomal dysfunction and oxidative stress in models of Snyder-Robinson syndrome. Nature Communications, 2017. 8(1).

57. Cosens, D.J. and A. Manning, Abnormal Electroretinogram from a Drosophila Mutant. Nature, 1969. 224(5216): p. 285–287.

